# Data requirements for accurate extinction-risk prediction in bistable populations

**DOI:** 10.64898/2026.06.19.733461

**Authors:** Adarshkrishnan Rajakumar, Pascal R. Buenzli, Matthew J. Simpson

## Abstract

Understanding and predicting extinction risk is a central challenge in population biology. Mathematical models incorporating Allee thresholds are commonly used to understand population dynamics and to assess extinction risks. Inaccurate predictions can have serious consequences for conservation management. In this simulation study, we develop a likelihood-based inference and prediction workflow to estimate parameters, including the Allee threshold and population diffusivity parameters, using noisy count data generated using a well-defined discrete model. Although parameters are identifiable according to commonly used criteria, the accuracy of resulting predictions depends strongly on the quantity, quality, collection time and spatial resolution of the data. Our workflow demonstrates that seemingly reliable parameter estimates can lead to inaccurate predictions, highlighting the need for careful consideration of data quality and quantity to guide extinction-risk modelling and prediction. Open source software is provided on GitHub to replicate and extend all results considered.

## 1 Introduction

Understanding the risk of population extinction is a central question in theoretical population biology that has long been explored using mathematical models [1–3]. Classical continuum models of population growth for a single species, such as the exponential and logistic growth models, cannot be used to study population extinction since the long-time solution of these models always predict population survival [1, 2]. This observation motivates the use of bistable equations since these models have the capacity to predict either survival or extinction, depending on whether the initial population density is greater than or less than some density threshold [2, 4]. These ideas are most simply encoded in the strong Allee effect model [5],

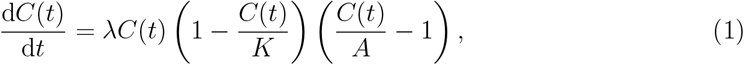

where *C*(*t*) ≥ 0 is the population density, *λ >* 0 is the growth rate, *K >* 0 is the carrying capacity density, and 0 *< A < K* is the Allee threshold density. Biologically, the Allee threshold implicitly models complex mechanisms such as difficulty in finding mates, reduced cooperative interactions, or increased susceptibility to predation at low population densities, which lead to negative net growth when the population falls below the Allee threshold density [2, 6, 7]. Solutions of Equation (1) have the property that *C*(*t*) → 0^+^ as *t* → ∞ provided *C*(0) *< A*, which models population extinction. Alternatively, *C*(*t*) → *K* as *t* → ∞ provided *C*(0) *> A*, which models population survival [2, 6]. Many generalisations of Equation (1) are possible, such as including diffusion-type transport operators to model spatial dispersion [1, 8, 9]. Regardless of these generalisations, a fundamental question remains about our ability to reliably estimate the Allee threshold *A* from data so that we can use mathematical models to make informed predictions about population survival and extinction.

Various approaches have been used to estimate Allee thresholds across a range of ecological applications, including populations of wolves [10], polar bears [11], gypsy moths [12], predator-prey systems [13, 14], and tumour cells [15]. Parameter estimation has also been performed using different types of data, such as pheromone trap records [16], time-series [17], and spatial data [18]. Relatively little attention has been paid to the identifiability of parameter estimates [19], or to understanding the data quality and quantity required for reliable extinction-risk predictions [20].

In this work, we explore the extent to which we are able to estimate parameters relevant to population survival and extinction using synthetic data generated by a discrete mathematical model. Working with a discrete model allows us to explore different types of realistic, noisy count data across a range of modelling scenarios. As we will explain, we use a continuum limit description of the discrete model which avoids concerns about model selection that inevitably arise when dealing with field or laboratory data [21, 22]. Using a discrete lattice-based model proposed by Li et al. [23], we generate realistic count data under situations leading to population survival or extinction. A key outcome of this work is the development and demonstration of an efficient likelihood-based workflow for parameter estimation [24, 25], identifiability analysis [26–29], and model-based prediction [30, 31] where we make use of efficient continuum-limit descriptions of the discrete model. A key motivation for this approach is that we avoid relying on expensive simulation-based inference methods since the discrete model is computationally demanding relative to the continuum-limit approximation.

The identifiability-estimation-prediction workflow is summarised in Figure 1. First, we generate spatial count data from the discrete stochastic model using equally spaced rectangular regions, which are often called *quadrats* [32–35] in the ecology literature, as shown in Figures 1(a)–(b). To describe this discrete count data using our continuum model, we evaluate the solution of the continuum limit model at the centre of each quadrat to give a prediction of the expected occupancy of each quadrat, as illustrated in Figures 1(c)–(d). This allows us to formulate a count–based likelihood function to quantify assess the likelihood of any particular parameterisation of the continuum model (e.g. ***θ*** = (*θ*_1_, *θ*_2_)^⊤^) [25]. We identify the maximum likelihood estimate (MLE) and construct associated parameter confidence sets, as shown in Figure 1(e). Finally, we use the parameter confidence sets to make quantitative predictions regarding the long-time survival or extinction of the population, as shown in Figure 1(f). As we will demonstrate and explore, our results illustrate that our ability to accurately forecast population survival or extinction depends strongly upon data quality and data quantity. In particular, we show that it is possible to arrive at identifiable and seemingly reliable parameter estimates that can lead to erroneous survival or extinction predictions. We explore the extent to which the initial spatial distribution of the population influences our ability to accurately forecast whether a population will become extinct. Open-source software is provided on GitHub [36] so that our results can be replicated and so that the workflow can be applied to different situations, such as working with populations involving different initial population sizes and spatial distributions.’

**Figure 1.**
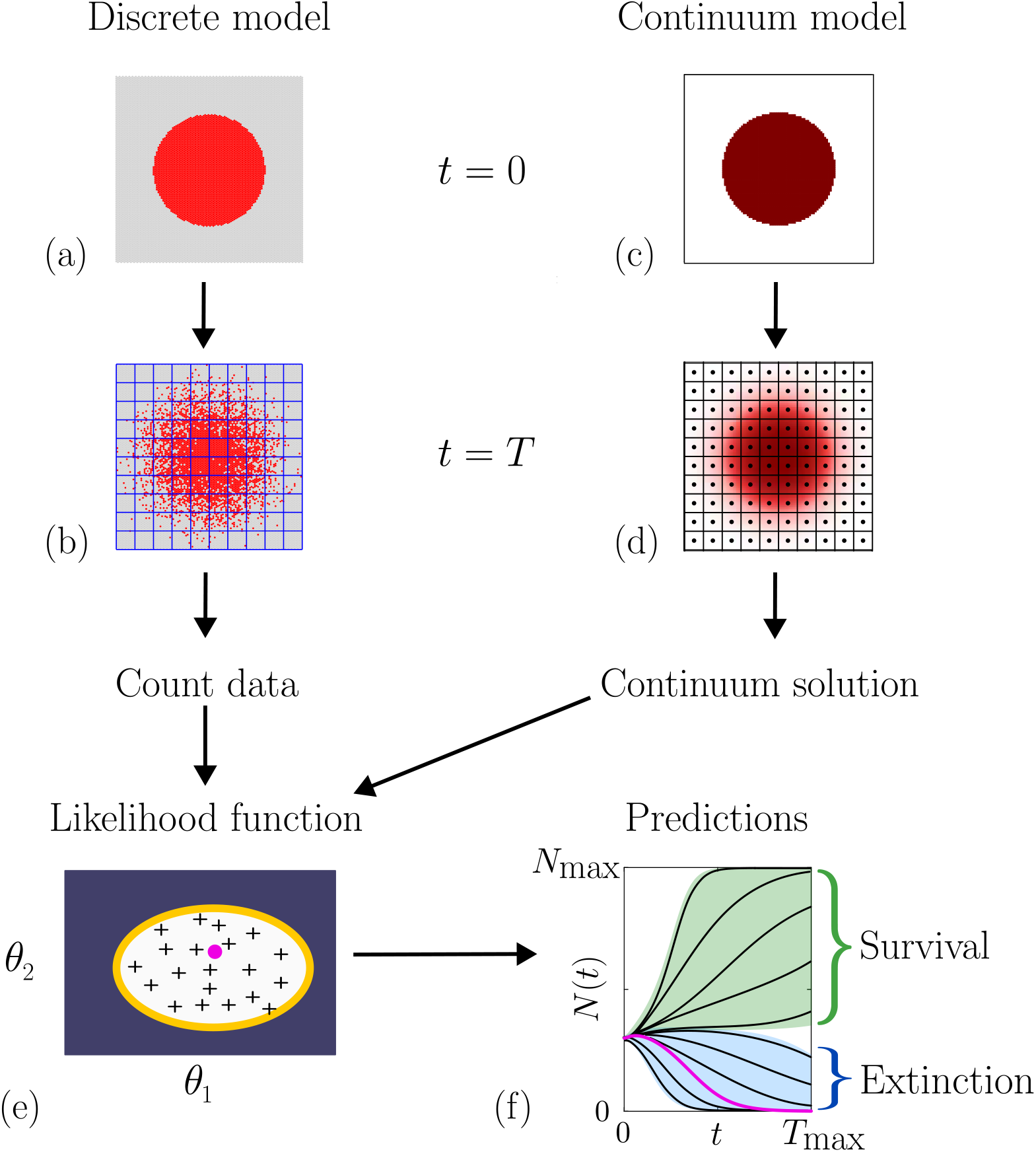
Workflow for identifiability, estimation and prediction: (a)–(b) Noisy count data from the discrete model at observation time *T* based on a grid of quadrats. (c)–(d) The solution of the continuum-limit description evaluated at the centre of each quadrat provides an estimate of the expected quadrat occupancy. (e)–(f) Count data and expected quadrat occupancy for parameters ***θ*** = (*θ*_1_, *θ*_2_)^⊤^ are used to evaluate a likelihood function, from which the MLE (purple dot) and parameter confidence set (yellow curve) are determined. Candidate parameter values (pluses) within the confidence set are used to construct prediction intervals shown in (f), where some trajectories indicate survival (green) and others indicate extinction (blue). MLE solution (solid purple curve) superimposed with other candidate solutions associated with parameters from within the prediction interval (solid black curves).

## 2 Mathematical models

To generate synthetic data, we use a stochastic hexagonal lattice-based discrete model with a bistable source term, as detailed in Section 2.1. This framework provides a controlled source of noisy, spatially structured count data in quadrats that incorporates the intrinsic stochasticity typical of realistic observations. By pairing stochastic count data with the continuum approximation discussed in Section 2.2, which captures the expected mean behaviour of the stochastic model, we perform likelihood-based parameter estimation, identifiability analysis and model-based prediction [25, 26]. Since the continuum model parameters corresponding to the simulation data are known, this approach allows us to rigorously evaluate how different data collection protocols affect parameter identifiability and the accuracy of population survival or extinction predictions.

### 2.1 Discrete mathematical model

We consider a two-dimensional lattice-based discrete model, where we use a hexagonal lattice with lattice spacing Δ *>* 0 as illustrated in Figure 2. This model was previously introduced and fully described by Li et al. [23], and so here we describe key features of the model only. Individuals are represented as agents that undergo crowding-dependent movement, proliferation, and death events as illustrated in Figure 2. We consider a hexagonal lattice with *I* × *J* lattice sites. Each site is indexed by **s** = (*i, j*), where *i* = 1, 2, 3, …, *I* and *j* = 1, 2, 3, …, *J*, and is associated with unique Cartesian coordinates given by

**Figure 2.**
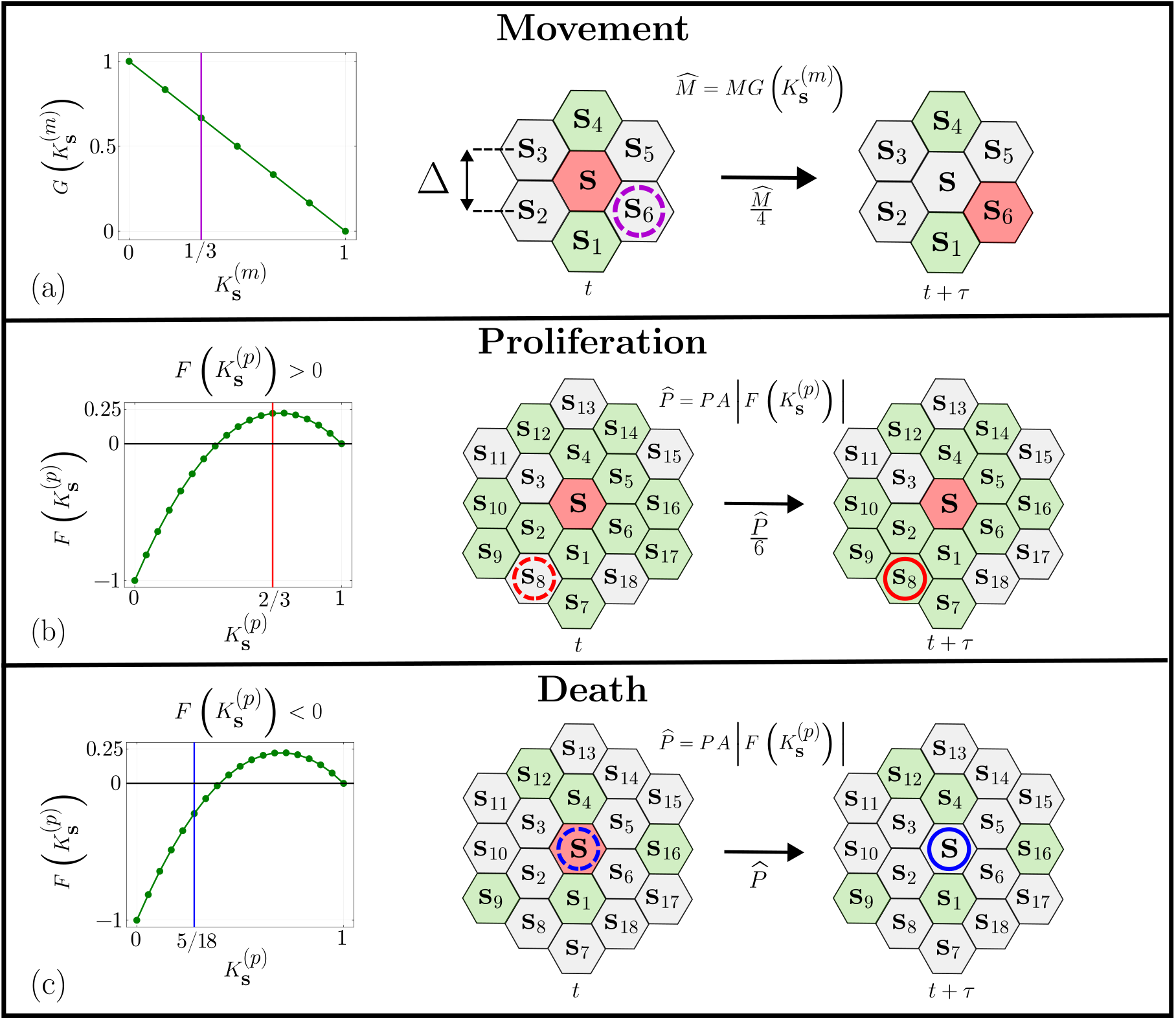
Crowding-dependent proliferation, death and movement in the discrete model. Several lattice fragments surrounding an occupied site **s** (red) are show, these lattice fragments include both occupied (green) and vacant sites (grey). In (a) the agent at site **s** moves randomly to one of the four neighbouring vacant sites (violet dashed) with probability 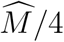, where 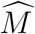 depends on the movement crowding function 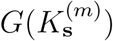 (green) evaluated at the local density (violet vertical line). In (b) we demonstrate a potential proliferation event and since 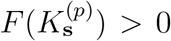 (green line) at the local density (vertical red line) the agent at **s** proliferates so that a new daughter agent is placed on a vacant site (red dashed) with probability 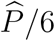. In (c) we demonstrate a potential proliferation event and since 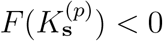 (green line) at the local density (vertical blue line) the agent at **s** dies with a probability 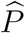.

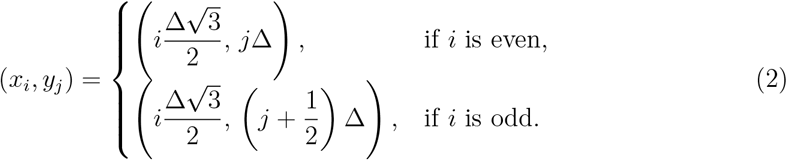

Simulations are performed using a discrete time framework where each time step has a constant duration *τ* . We work with dimensionless simulations by setting Δ = *τ* = 1, and noting that physical units can be recovered by appropriate rescaling of Δ and *τ* [25]. At time *t*, each site **s** is either occupied *C*_**s**_(*t*) = 1, or vacant *C*_**s**_(*t*) = 0, with at most one agent per site, which means that the discrete model is closely related to an exclusion process. For convenience we denote the occupancy of each lattice site using *C*_**s**_(*t*) and *C*_*i, j*_(*t*) interchangeably. In Cartesian coordinates, the domain has width 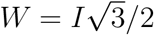 and height *H* = *J* when Δ = 1.

In the discrete model agents undergo motility and proliferation events that are simulated using the random sequential update algorithm [37]. If there are *N* (*t*) agents on the lattice at time *t* we randomly select *N* (*t*) agents, one at a time with replacement, and give those agents an opportunity to move. After these *N* (*t*) potential movement events have been assessed, we randomly select *N* (*t*) agents, one at a time with replacement, and allow each of these to attempt to proliferate. The outcome of these potential motility and proliferation events depends upon the local crowding conditions, as we now explain.

To incorporate crowding in the motility mechanism, we introduce the quantity 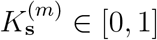, which is a non-dimensional measure of the density surrounding site **s** and is given by

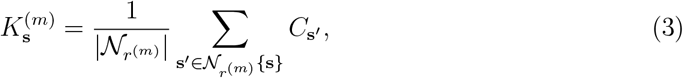

where, *r*^(*m*)^ = 1, 2, 3, … is an integer number of concentric rings of sites centred at site **s**, and 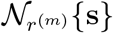 is the collection of sites contained within the movement template. The movement template is taken to be all sites contained within a concentric region about site **s**, noting that the total number of sites within the template of radius 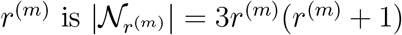 [38], as shown in Figure 2. When an agent at a site **s** is selected for a potential motility event, that agent will attempt to move to a vacant site in 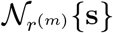 with probability 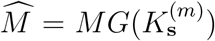, where 0 ≤ *M* ≤ 1 denotes the probability that an isolated agent moves during a time interval of duration *τ* . Here, 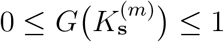 is the *movement crowding function* that describes how local density impacts agent motility. Throughout this work we choose 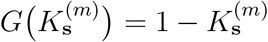 so that *G*(1) = 0 and *G*(0) = 1. This choice reflects the fact that the motility of isolated agents with *G*(0) = 1 are not impacted by crowding, whereas agents packed at the maximum density with *G*(1) = 0 are unable to move owing to crowding. If the agent moves, it relocates from **s** to a randomly chosen vacant site within 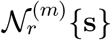 so that the probability that an agent located at site **s** moves into any particular vacant site is 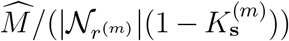. Figure 2(a) illustrates the movement mechanism with 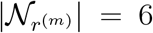. In this case, the agent at site **s** has two occupied neighbouring sites, giving 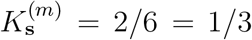. The probability that this agent moves is therefore 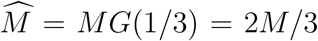. Since four sites in 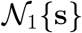 are vacant, the probability of moving to any one of these vacant sites is 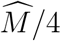.

In the discrete model we make a distinction between the template sizes for motility and proliferation by introducing *r*^(*p*)^ = 1, 2, 3, …, as the template size for proliferation. Using the same approach for modelling crowding in the motility mechanism, we work with a local density 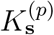, where 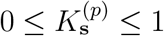, to account for crowding in proliferation events. All results in this work correspond to *r*^(*m*)^ = 1 and *r*^(*p*)^ = 4 to be consistent with Li et al. [23]. For each prospective proliferation (or death) event, an agent at site **s** attempts to proliferate (or die) with probability 0 ≤ *P* ≤ 1 within each time step of duration *τ* . We set 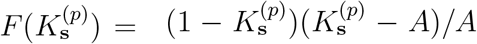, which ensures that *F* (0) = −1 and *F* (1) = 0, where 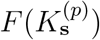 is the *proliferation crowding function*. Potential proliferation events succeed with probability 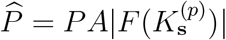. The proliferation crowding function describes how local crowding around **s** influences whether agents proliferate if 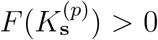, or die if 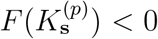. As we discuss in Subsection 2.2 our definition of 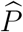 ensures that the source term in the continuum limit description is independent of *A*. Our choice of 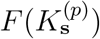 with 0 *< A <* 1, yields a cubic source term in the continuum limit with strong Allee-type kinetics, where *A* is the Allee threshold [6, 39, 40]. For all results in this work, we set *A* = 4 × 10^−1^ in the discrete model. For a proliferation event, a daughter agent is placed on a randomly chosen vacant site within 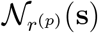, and during a death event the agent at **s** is removed. At the end of each time step, after all *N* (*t*) agents on the lattice have been considered for potential proliferation (or death), we update *N* (*t*+*τ*) accordingly. Figure 2(b)–(c) depicts this proliferation mechanism with *r*^(*p*)^ = 2. In Figure 2(b), there are five neighbouring agents giving 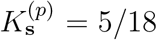. Since *F* (5*/*18) *<* 0 the agent at **s** is removed with probability 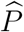. In Figure 2(c), the agent at **s** has 12 neighbours giving 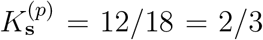. The probability of proliferation is *PA*|*F* (2*/*3)|. Since *F* (2*/*3) *>* 0 and there are six vacant sites in N_2_{**s**}, a new agent is placed on one of these with probability 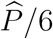.

All discrete simulations are conducted with *M* = 1, *P* = 2.5 ×10^−3^, *r*^(*m*)^ = 1 and *r*^(*p*)^ = 4, on a lattice of dimensions *I* × *J* = 116 × 100, corresponding to a physical domain of *W* × *H* ≈ 100 × 100.

### 2.2 Continuum mathematical model

The mean behaviour of the discrete model is described by a partial differential equation (PDE) that was derived by Li et al. [23]. Working with a continuum approximation provides us with a substantial computational advantage since the approximate continuum PDE can be solved very efficiently compared to the computational time required to simulate the discrete model. Moreover, continuum models can provide analytical insight into the population dynamics rather than relying on discrete simulations only. The continuum limit PDE is given by [23]

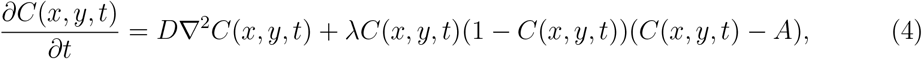

*C*(*x, y, t*) ≤ 1, and *A* is the Allee threshold with 0 *< A <* 1, corresponding to the parameter *A* in the proliferation probability 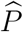 of the discrete model. Furthermore, we have

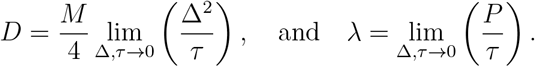

Here, *D >* 0 is the macroscopic diffusivity obtained in the limit of Δ → 0 and *τ* → 0 while keeping Δ^2^*/τ* to be O(1) [41, 42] and *λ* is the proliferation rate. The source term in Equation (1) is a cubic bistable model with a carrying capacity of one. This feature arises because the maximum number of agents per lattice site in the discrete model is one, implying that 0 *< A <* 1. To obtain a well-defined continuum limit model, we require *λ* to be O(1) which introduces the restriction that *P* is O(*τ*) [43] in the limit that *τ* → 0. This means that we expect the continuum limit to be accurate for *P/M* ≪ 1. Since Δ = *τ* = 1 in the discrete model, we have *D* = *M/*4 and *λ* = *P* in the continuum formulation. Henceforth, we set *A* = 4 × 10^−1^ and *D* = 2.5 × 10^−1^ in the continuum model as *reference* parameters that are equivalent to *A* = 4 × 10^−1^ and *M* = 1 in the discrete model with Δ = *τ* = 1. We treat these reference parameters as the ground truth for identifiability, inference and prediction. In Section 4, we perform rigorous checks to ensure that the continuum model provides an excellent approximation to the mean behaviour of the data collected from the discrete model.

The diffusion term in Equation (1) is linear. This follows from the choice of the movement crowding function in the discrete model, 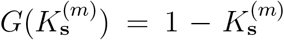, which yields a constant macroscopic diffusivity in the continuum limit [44]. Alternative crowding functions lead to density-dependent non-linear diffusion terms [44] that we do not consider here.

An important point to note is that many continuum mathematical models used to describe population biology phenomena involve ODE and PDE descriptions that are similar to Equation (1) in the sense that the dependent variable provides a measure of population density [1– 3]. In contrast, most data describing population biology experiments involve non-negative, integer-based counts of individuals [31]. Key aspects of the novelty of our workflow involve maintaining the computational efficiency of ODE and PDE–based models in a framework that keeps these modelling frameworks relevant to working with count-based data. This is a subtle but important point that is often overlooked in ODE and PDE-based modelling [31].

## 3 Quantifying count data

In this Section, we describe how we generate count data from the discrete model under four different initial conditions. For each discrete simulation we specify an initial distribution of agents given by

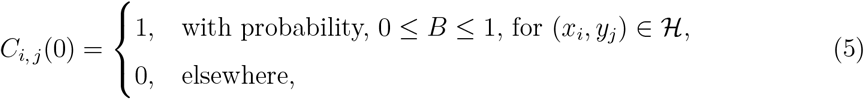

where ℋ denotes the initially occupied region in Cartesian space, with (*x*_*i*_, *y*_*j*_) defined previously in Equation (1). The analogous initial condition for the continuum model is

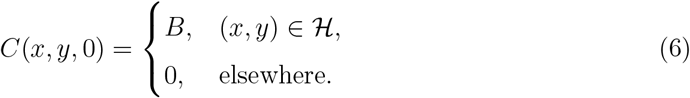

By varying our choice of ℋ and *B*, we generate four distinct initial conditions: (i) a well-mixed initial condition shown in Figure 3(a); (ii) a one-dimensional (1D) column initial condition shown in Figure 3(b); (iii) a two-dimensional (2D) square initial condition shown in Figure 3(c); and, (iv) a 2D circular initial condition shown in Figure 3(d).

**Figure 3.**
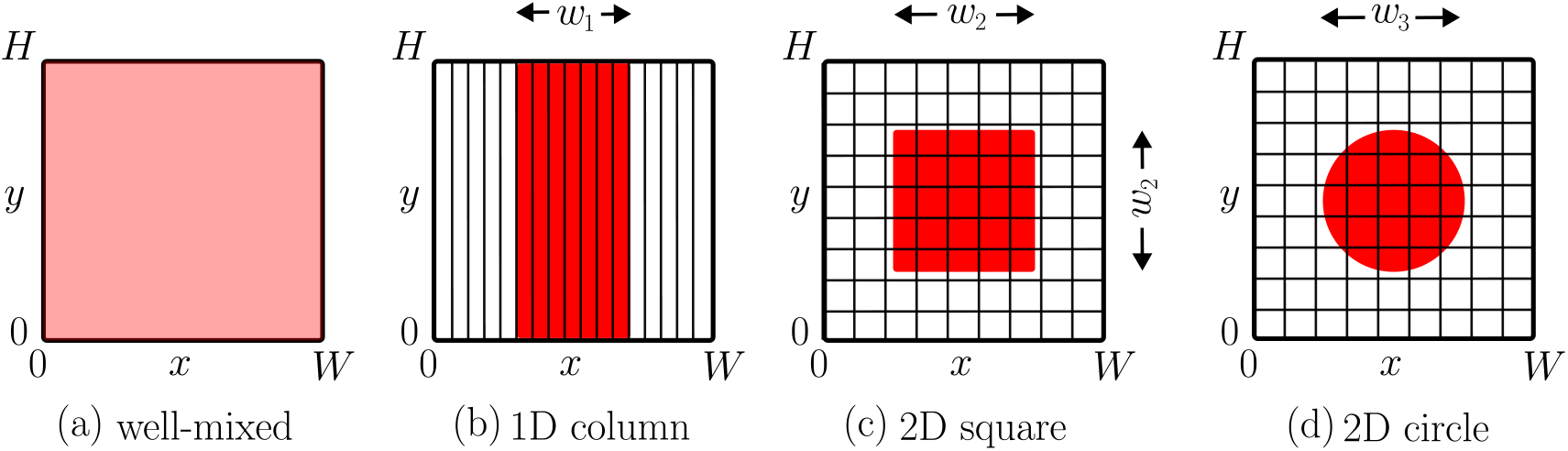
Quadrat partitioning. (a) Well-mixed distribution, where the entire domain is counted within a single quadrat, *U* = *V* = 1. (b) 1D column initial distribution of width *w*_1_, partitioned into vertical strip-shaped quadrats with *V* = 1 and *u* = 1, 2, 3, …, *U* . (c) 2D square initial distribution of width *w*_2_ partitioned into rectangular quadrats with *u* = 1, 2, 3, …, *U* and *v* = 1, 2, 3, …, *V* . (d) 2D circular initial distribution of diameter *w*_3_ partitioned into rectangular quadrats with *u* = 1, 2, 3, …, *U* and *v* = 1, 2, 3, …, *V* . Note that we specify *w*_1_, *w*_2_, and *w*_3_ as physical lengths.

To quantify the spatially structured count data generated by the discrete model simulation, we partition the *H* × *W* domain into a coarser discretisation of *U* × *V* equally spaced rectangular–shaped quadrats. This approach models ecological sampling methods, where quadrats are traditionally used to count spatially structured populations [32]. We define quadrats in continuous space, with each quadrat indexed by (*u, v*), where *u* = 1, 2, 3, …, *U* and *v* = 1, 2, 3, …, *V* . Values of *U* and *V* are chosen depending on the initial condition, as illustrated in Figure 3. Each quadrat has a width of *W/U* and a height of *H/V* . At any time *t*, we record the number of agents, *N*_*u,v*_(*t*), whose Cartesian coordinates (*x*_*i*_, *y*_*j*_) lie within the boundaries of quadrat (*u, v*). We assemble these counts across all quadrats into an observation vector of length *UV*, which we denote as **N**^obs^(*t*). When partitioning an *I* × *J* discrete lattice using *U* × *V* quadrats, the average number of lattice sites per quadrat is (*IJ*)*/*(*UV*), however either *I/U* or *J/V* are not integers, then rounding impacts the actual number of lattice sites within a quadrat. We denote the total number of lattice sites in a quadrat indexed (*u, v*) as 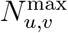, which is bounded above by a theoretical maximum 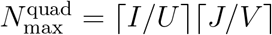, so that 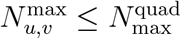 for all (*u, v*).

Our approach to partitioning the spatial region into quadrats is tailored to match the symmetry of each initial spatial distribution of agents as follows:

1. The well-mixed initial distribution is treated as a single quadrat (*U* = *V* = 1) as illustrated in Figure 3(a) where **N**^obs^(*t*) has a single element corresponding to the total population, *N* (*t*). The total number of lattice sites in the quadrat is 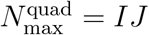.
2. For the 1D column initial distribution, we use *u* = 1, 2, 3, …, *U* equally spaced vertical strips with *V* = 1, as illustrated in Figure 3(b). Each strip has a width of *W/U* and height of *H*, with 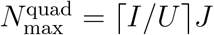.
3. The 2D square initial distribution and the 2D circular initial distribution use a regular grid with *u* = 1, 2, 3, …, *U*, and *v* = 1, 2, 3, …, *V* giving *UV* rectangular-shaped quadrats, as illustrated in Figure 3(c)–(d). Each quadrat has a width *W/U* and height *H/V*, with 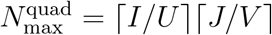.

We compare discrete counts *N*_*u,v*_(*t*) within each quadrat, where 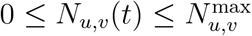, to the corresponding quantities obtained from solving the continuum model by defining

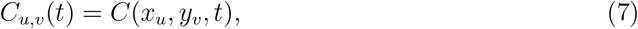

where (*x*_*u*_, *y*_*v*_) is the centre of the quadrat indexed by (*u, v*). The continuum analogue of *N*_*u,v*_(*t*) is

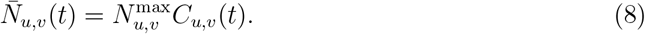

While quadrat counts from the discrete simulations inherently include stochastic fluctuations, the continuum model gives a deterministic, noise-free solution. In Section 4 we will compare quadrat-based count data using both the discrete and continuum modelling approaches.

## 4 Relating continuum and discrete models

In this Section, we present a suite of discrete simulations to demonstrate that we can use Equation (1) to accurately approximate count data from the discrete model under a range of conditions. Utilising the quadrat partitioning framework established in Section 3, we show that the continuum model provides a good approximation for both quadrat counts *N*_*u,v*_(*t*), and the total population size *N* (*t*).

The total number of agents in the discrete simulation at time *t* is given by

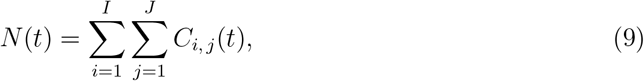

which is a bounded non-negative integer, 0 ≤ *N* (*t*) ≤ *IJ* . The corresponding continuum quantity 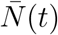 is obtained by solving Equation (1) numerically (see Appendix A) to give *C*(*x, y, t*), from which we have

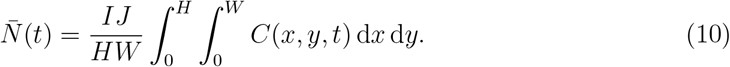

The factor (*IJ*)*/*(*HW*) in Equation (1) is the number of lattice sites per unit area, which simplifies to 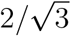 in our case when Δ = 1. We approximate the integral in Equation (1) using the rectangle rule, and we will comment on the appropriateness of this choice later. It is important to note that while 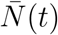 provides a summary of the total population dynamics which ignores the spatial distribution of agents on the lattice, the parameter estimation approach described in Section 5 relies on the spatially structured quadrat counts *N*_*u,v*_(*t*).

For all initial configurations, we simulate both an extinction scenario and a survival scenario by adjusting *B* and ℋ to vary *N* (0), while keeping all other parameters in the discrete model fixed. Results in Figure 4 correspond to a well-mixed initial condition where agents are uniformly distributed across the entire domain. Simulation snapshots are presented in Figure 4(a)–(c) where we see that the population eventually becomes extinct, whereas similar snapshots in Figure 4(e)–(g) show that the population eventually survives. For the well-mixed case the macroscopic spatial density gradient vanishes everywhere so that ∇*C* = **0** and *C*(*x, y, t*) = *C*(*t*). Under these conditions, the PDE given by Equation (1) reduces to the ordinary differential equation (ODE) [43]

**Figure 4.**
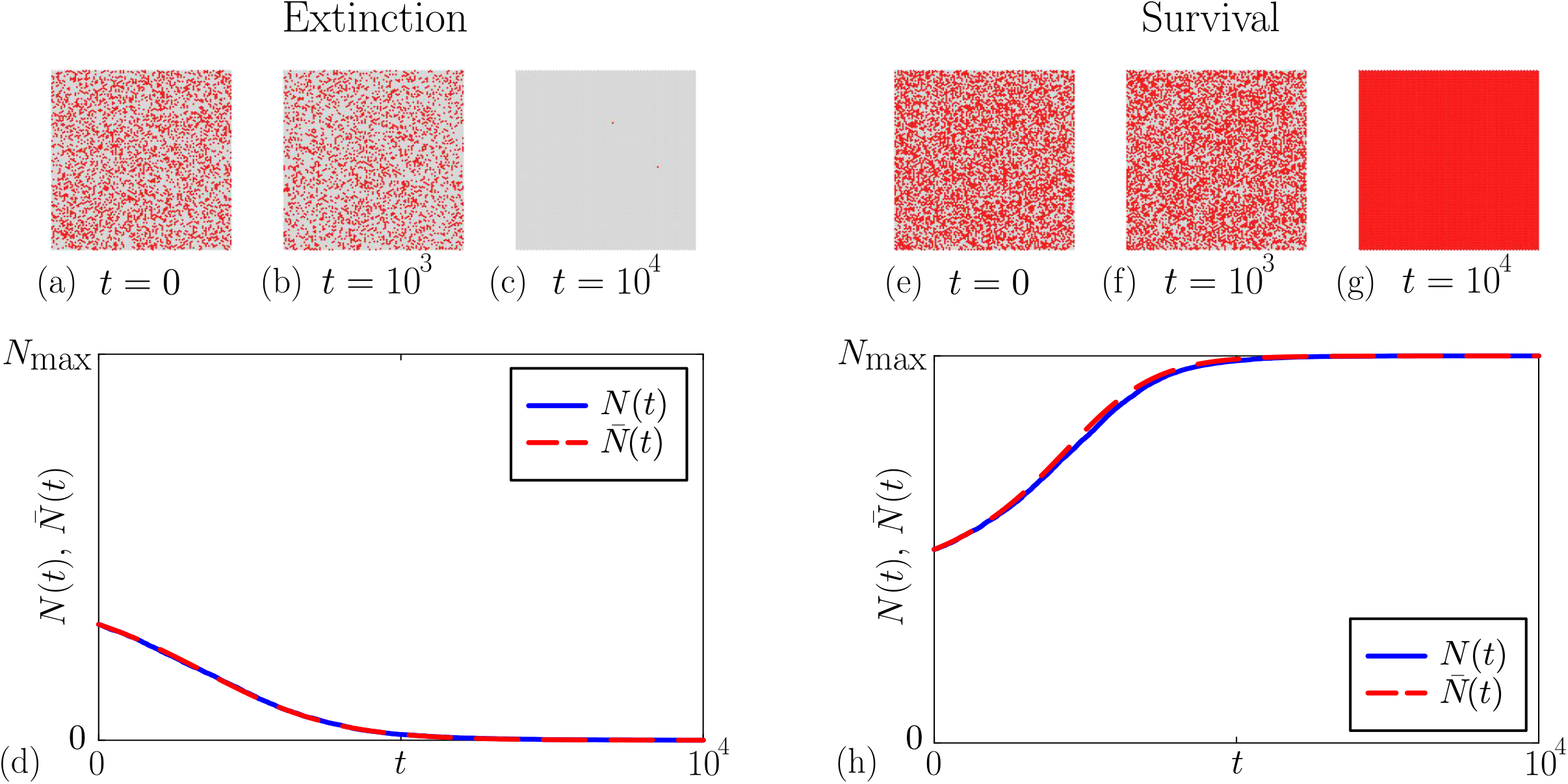
Discrete simulation data and the solution of the continuum limit model for the well-mixed initial condition. (a) Initial lattice configuration for the extinction scenario with *B* = 3 × 10^−1^. (b)–(c) Snapshots at *t* = 10^3^ and 10^4^. (e) Initial lattice configuration for the survival scenario with *B* = 5 × 10^−1^. (f)–(g) Snapshots at *t* = 10^3^ and 10^4^. (d) and (h) compare *N* (*t*) (blue) and 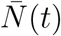 (red dashed) for the extinction and survival scenarios, respectively, with 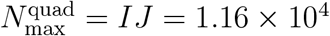. Simulation parameters are *M* = 1, *A* = 4 × 10^−1^, *P* = 2.5 × 10^−3^, *r*^(*m*)^ = 1 and *r*^(*p*)^ = 4.

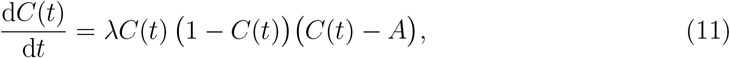

where 0 ≤ *C*(*t*) ≤ 1 is the spatially invariant average agent occupancy at time *t*. To use the continuum limit model, we solve Equation (1) numerically (see Appendix A) to give estimates of *C*(*t*). In the well-mixed scenario, we collect count data using a single quadrat spanning the entire lattice (*U* = *V* = 1). Consequently, *N*_1,1_(*t*) = *N* (*t*) and the expected count is 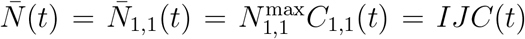. Results in Figure 4(d) and Figure 4(h) compare *N* (*t*) and 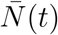 for both the extinction and survival cases, respectively. These comparisons confirm that the solution of the continuum limit model provides an excellent approximation of the discrete model.

We now consider a different initial configuration in Figure 5 where we present snapshots from the discrete model where agents are initially placed in a column-shaped initial arrangement. In these simulations agents are placed at maximal density *B* = 1, within a vertical strip of width *w*_1_. Results in Figure 5(a)–(c) show snapshots from the discrete model where the initial column width is sufficiently narrow that the population eventually goes extinct. In contrast, results in Figure 5(f)–(h) show snapshots from the discrete model with the same parameters except that the initial column width is sufficiently wide that the population eventually survives.

**Figure 5.**
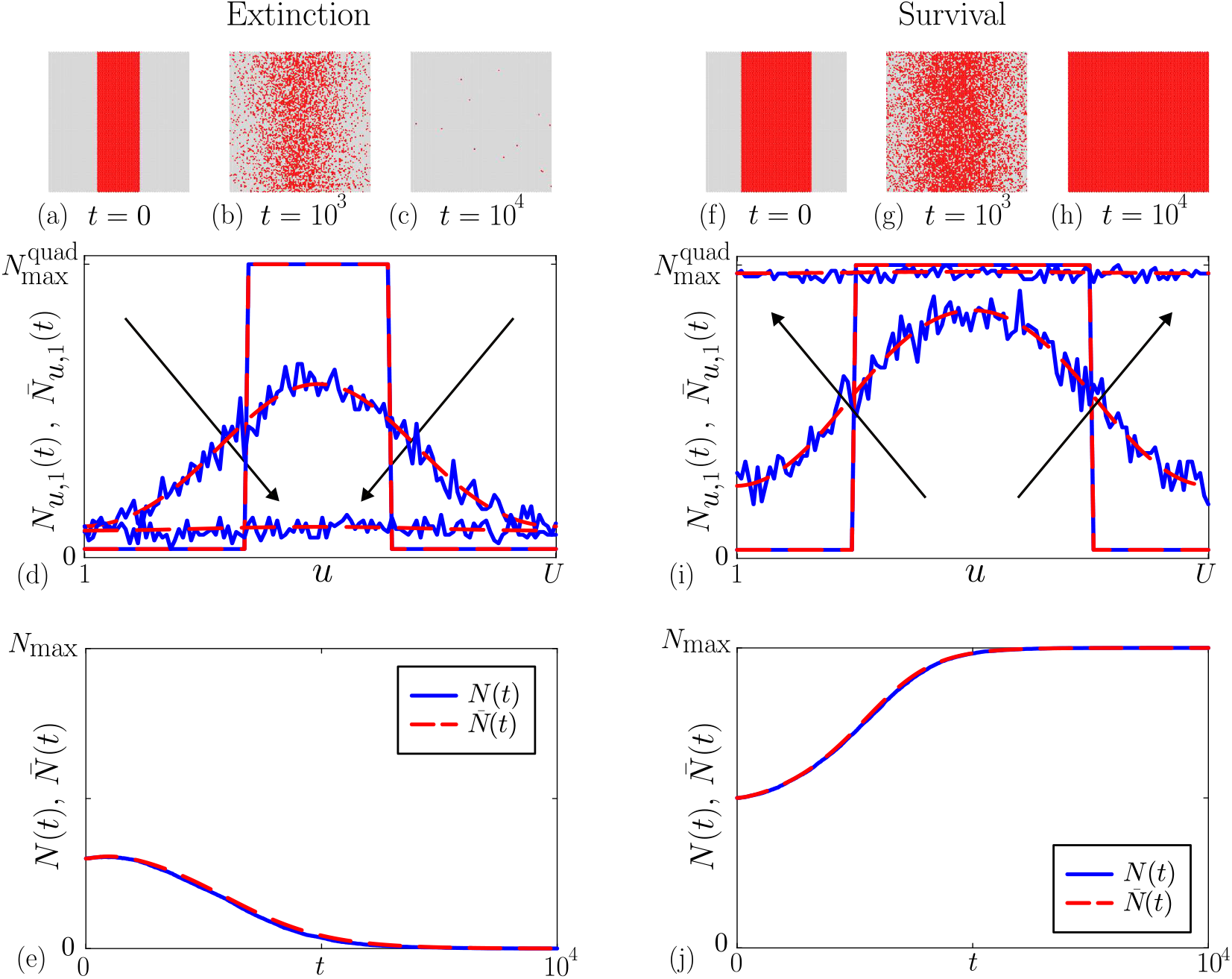
Discrete simulation data and the solution of the continuum limit model for the 1D column initial condition. (a) Initial lattice configuration with column width *w*_1_ = 30. (b)–(c) Simulation snapshots at *t* = 10^3^ and 10^4^. (d) Comparison of *N*_*u*, 1_(*t*) (blue) and 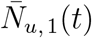 (red dashed) at *T* = 0, 10^3^ and 5 × 10^3^ where the arrows show the direction of increasing time, and 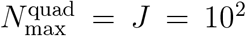. (e) Comparison of *N* (*t*) (blue) and 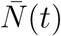 (red dashed) with *N*_max_ = *IJ* = 1.16 × 10^4^. (f) Initial lattice configuration where column width *w*_1_ = 50. (g)–(h) Simulation snapshots at *t* = 10^3^ and 10^4^. (i) Comparison of *N*_*u*, 1_(*t*) (blue) and 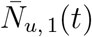 (red dashed) where the arrows show the direction of increasing time. (j) Comparison of *N* (*t*) (blue) and 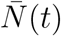 (red dashed). Simulation parameters are *M* = 1, *A* = 4 × 10^−1^, *P* = 2.5 × 10^−3^, *r*^(*m*)^ = 1 and *r*^(*p*)^ = 4.

To describe these simulations using the continuum limit description, we note that the macroscopic density of agents is independent of vertical position [43]. Under these conditions *C*(*x, y, t*) = *C*(*x, t*), and Equation (1) simplifies to

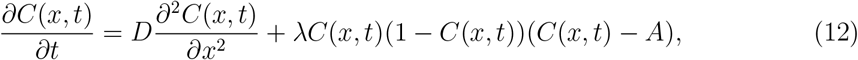

where 0 ≤ *C*(*x, t*) ≤ 1, is the average agent occupancy at position 0 *< x < W* and time *t >* 0. To proceed, we solve Equation (1) numerically (see Appendix A) using periodic boundary conditions and an initial condition given by Equation (1) to give *C*(*x, t*). To quantify the spatial count data, we partition the domain into vertical column–shaped quadrats by setting *u* = 1, 2, 3, …, *U* = *I* and *V* = 1. In this case we make the simple choice of setting *U* = *I* which ensures that each column of the hexagonal lattice maps uniquely to a single quadrat of width 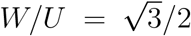. Alternative choices of *U* could also be considered within this framework. We compute the counts of agents in each quadrat using the discrete simulation data as

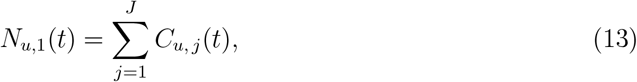

where each *u* corresponds to a column index *i*. Because every column contains exactly *J* lattice sites we have 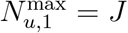 for *u* = 1, 2, 3, …, *U* . Consequently, the continuum analogue of these column counts is 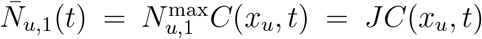. This expression can be evaluated with the numerical solution of Equation (1).

The absence of spatial gradients in the *y*-direction means that Equation (1) simplifies to

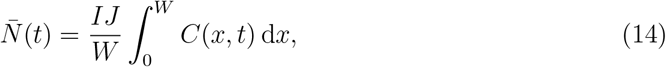

which we estimate using the rectangle rule by discretising the domain 0 *< x < W* with 300 equally spaced mesh points.

Results in Figure 5(d) and Figure 5(i) compare *N*_*u*,1_(*t*) and 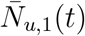 for the extinction and survival scenarios, respectively. This comparison shows that 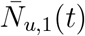 gives an excellent approximation of the mean behaviour of *N*_*u*,1_(*t*), but without capturing the fluctuations in the column counts. Results in Figure 5(e) and Figure 5(j) show the evolution of the total discrete population *N* (*t*) for the extinction and survival scenarios, respectively. Visual comparison of *N* (*t*) and 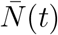 in Figure 5(e) and Figure 5(j) shows that the continuum model gives an excellent approximation of the discrete data, indicating that our use of the simple rectangle rule for quadrature performs well for our purpose.

Finally, we consider a third initial configuration in Figure 6 that involves initialising agents in a circular region so that lattice sites within a particular circular region are initialised at maximal density, *B* = 1. Results in Figure 6(a) show snapshots of the discrete model where the radius of the initially occupied region is 30 and we see that the population eventually goes extinct. Results in Figure 6(d) show snapshots from the discrete model with the same parameters except that the radius of the initially occupied region is now 40 and we see that the population survives in these simulations. The corresponding solution of Equation (1) is shown in Figure 6(b) for the extinction scenario, and in Figure 6(g) for the survival scenario. These simple visualisations indicate that the solution of the continuum model provides a good match with the discrete data. In Figure 6(c)–(d) and Figure 6(h)–(i) we provide a visual comparison of quadrat-based counts from the discrete model *N*_*u,v*_(*t*) with the continuum analogue 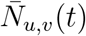, confirming a good match between the discrete and continuum quantities with the continuum model predicting the average behaviour of the noisy count data.

**Figure 6.**
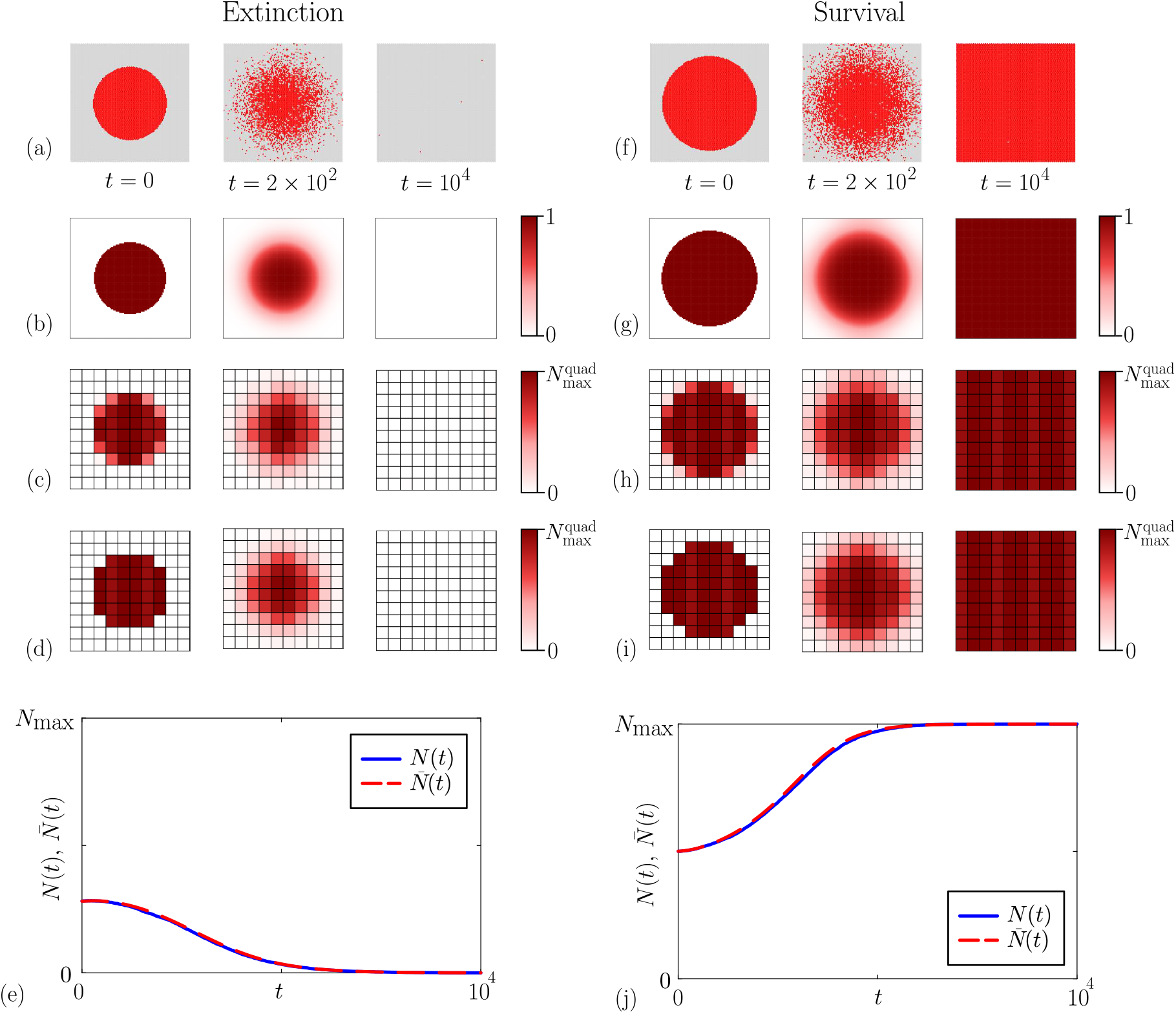
Discrete simulation data and the appropriate solution of the continuum limit model for the 2D circular initial condition. (a) Discrete simulation snapshots where the initially occupied region is a circle of radius 30 and the population becomes extinct. (b) The numerical solution of Equation (1), *C*(*x, y, t*) at *t* = 0, 10^2^ and 10^4^. (c) Spatial distribution of quadrat counts *N*_*u,v*_(*t*), the colour of each quadrat indicates the number of agents within that quadrat. The colour bar has 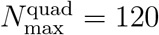. (d) Spatial distribution of the continuum quadrat count, 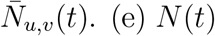 (blue) and 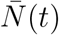 (red dashed) with *N*_max_ = *IJ* = 1.16 × 10^4^. (f) Discrete simulation snapshots where the initially occupied region is a circle of radius 40 and the population survives. (g) The numerical solution of Equation (1), *C*(*x, y, t*) at *t* = 0, 10^2^ and 10^4^. (h) Spatial distribution of *N*_*u,v*_(*t*). (i) Spatial distribution of 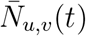. (j) *N* (*t*) (blue) and 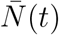 (red dashed). Simulation parameters are *M* = 1, *A* = 4 × 10^−1^, *P* = 2.5 × 10^−3^, *r*^(*m*)^ = 1 and *r*^(*p*)^ = 4.

Interestingly, there is a small discrepancy between *N*_*u,v*_(0) and 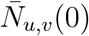 in Figure 6(c)–(d) and Figure 6(h)–(i), specifically for quadrats located near the edge of the initial distribution. This discrepancy arises because the continuum approximation is evaluated strictly at the centre of each quadrat. For partially occupied quadrats, our approach overestimates the occupancy if the centre of the quadrat lies within ℋ, or underestimates the occupancy if the centre of the quadrat lies outside ℋ. Comparing *N* (*t*) and 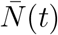 in Figure 6(e) and Figure 6(j) demonstrate strong agreement between the continuum approximation and discrete simulation data. Here, 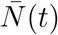 is calculated using Equation (1) by applying the rectangle rule on a uniform 100 × 100 mesh. The agreement between *N* (*t*) and 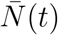 indicates that our simple approach to quadrature is appropriate, and the small discrepancies in quadrat-based counts at *t* = 0 are not important when comparing the evolution of the total population across the entire lattice.

## 5 Likelihood-based estimation and identifiability

Our results in Section 4 confirm that the solution of the continuum limit model can be used to accurately predict count data from the discrete model for different initial conditions, including both extinction and survival scenarios.

In this Section, we outline the methods used to estimate the parameters *D* and *A* from synthetic data generated by a single realisation of the discrete model where data are reported in terms of quadrat-based noisy count data. We focus on estimating parameters *D* and *A* using the corresponding continuum limit description while keeping *λ* and *C*(*x, y*, 0) fixed. When using the continuum approximation, this initial occupancy is *C*(0) for the well-mixed initial condition, *C*(*x*, 0) for the 1D column initial condition, and *C*(*x, y*, 0) for the 2D square and circular initial conditions. These differences in the shapes of the initial populations are matches with our choice of quadrat shape. This approach allows us to evaluate how accurately *D* and *A* are recovered across different spatial configurations.

The four different initial conditions we consider can be classified in two broad classes of inference problems. For data generated from the well-mixed initial condition, the continuum limit description, Equation (1) is independent of *D* so we aim to infer *A* only. This means that our unknown parameter vector has a single component, ***θ*** = (*A*)^⊤^. For data generated from the 1D column and 2D initial configurations, the continuum limit descriptions (Equation (1) and Equation (1)) involves both *A* and *D* so we target two parameters, ***θ*** = (*A, D*)^⊤^.

For a collection of count data in various quadrats, the number of agents in each quadrat is denoted by *N*_*u,v*_(*t*) and the total number of lattice sites in each quadrat is denoted by 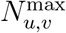. For bounded integer data, it is natural to assume that the probability of observing *N*_*u,v*_(*t*) counts in a quadrat (*u, v*) follows a binomial distribution with 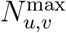 trials and a success probability *C*_*u,v*_(*t*) ∈ [0, 1] [25, 31]. This assumption leads to a log-likelihood function

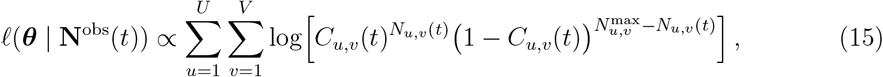

where we evaluate *ℓ* by setting the proportionality constant to unity [31]. Working with the binomial log-likelihood function is important because this captures the discrete nature of count data, and respects the fact that counts are non-negative and bounded from above. Unlike the more standard approach of assuming that data are normally distributed about the solution of some mathematical model [31, 45], working with the binomial distribution is attractive since there are no additional parameters in the noise model to estimate.

Given our likelihood function, the maximum likelihood estimate (MLE) is the best-fit parameter estimate [26, 46], defined as

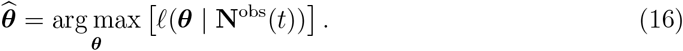

We evaluate this optimisation by discretising the parameter space and searching across a uniform grid of discrete *ℓ* values to find ***θ*** which maximises *ℓ* [47]. For problems involving a well-mixed initial condition, the parameter space involves only one unknown *A*, and we simply discretise the interval *A*^−^ ≤ *A* ≤ *A*^+^ uniformly with 300 mesh points. For problems involving the 1D column, 2D square and 2D circular initial conditions, the parameter space involves both *A* and *D* and we uniformly discretise the region defined by *A*^−^ ≤ *A* ≤ *A*^+^ and *D*^−^ ≤ *D* ≤ *D*^+^ using 100 × 100 mesh points.

Given the MLE, the normalised log-likelihood is

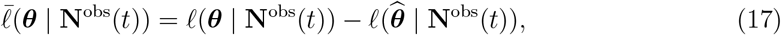

so that 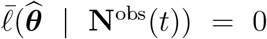. Working with the normalised log-likelihood allows us to construct asymptotic parameter confidence sets using Wilks’ theorem which states that the likelihood-ratio statistic converges in distribution to a *χ*^2^ variable in the large data-limit [24, 26, 48]. This allows us to define a likelihood-based threshold 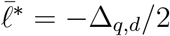, where Δ_*q,d*_ is the *q*th quantile of the *χ*^2^ distribution with *d* degrees of freedom [26, 48, 49], where *d* is the number of free parameters. Here we will work with the 0.95 quantile, giving 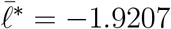 for problems involving one free parameter, and 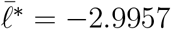 for problems involving two free parameters. With these ingredients we may then determine various confidence sets in the parameter space by identifying the region in the parameter space where 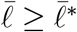. If this region is confined to a relatively small region of the parameter space, then the model parameters are practically identifiable from the data, whereas if the confidence set is unbounded or extends to the boundaries of the parameter space, the problem is poorly identifiable [26, 27].

To examine how uncertainty in parameter values within the confidence set propagate into variability in the predicted population dynamics, we sample values of ***θ*** from within the parameter confidence set using rejection sampling to generate *S* parameter samples [26, 31]. For each of the *S* parameter combinations we use the appropriate continuum approximation and Equation (1) to obtain *S* continuous predictions of the population dynamics, 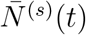 for *s* = 1, 2, 3, …, *S* [26, 31]. Each of these continuous predictions can be interpreted as a mean trajectory that neglects any stochastic variability [26, 31]. To approximately describe the variability about each mean trajectory, we use the binomial noise model to quantify fluctuations in *N* (*t*) about the mean. A standard approach to quantifying the width of a probability distribution is to compute various quantiles [26]. We compute the 2.5% and 97.5% quantiles of the binomial noise model at discrete time points *t*_*k*_ = *δt*(*k*−1) for *k* = 1, 2, 3, …,. Evaluating these quantiles for a specific parameter sample *s* at the *k*th time point yields an interval about the mean trajectory, 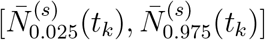 [26]. This interval provides a simple way to model variability implied by the noise model in terms of the 95% prediction interval at this time for that particular parameter realisation. Taking the union of such intervals at time *t*_*k*_ across all *s* parameter samples gives 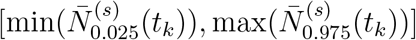 for *k* = 1, 2, 3, …, which we report here as an approximate 95% prediction interval for data realisations [26, 31]. In this work we refer to these intervals as *prediction intervals*, however strictly speaking, they are tolerance intervals as they incorporate both the uncertainty in estimating the model parameters and variability in future data realisations [31, 50].

## 6 Results and Discussion

Using count data from discrete simulations and the corresponding continuum analogues, we now implement the identifiability, estimation and prediction workflow to assess predictive accuracy across different initial conditions, quadrat sampling strategies, and observation times.

### 6.1 Well-mixed initial condition

First, we consider a well mixed initial distribution of agents and estimate ***θ*** = (*A*)^⊤^. We take the parameter space to be in the range 2 × 10^−1^ ≤ *A* ≤ 6 × 10^−1^, our results (below) confirm that this choice of parameter space is appropriate. Results in Figure 7(a) show a series of snapshots of a discrete simulation with *A* = 4 × 10^−1^ in which the population eventually goes extinct. Using count data at time *T* = 1 × 10^2^, we obtain 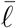 shown in the upper panel of Figure 7(c). The MLE is 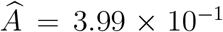, which is extremely close to the expected result *A* = 4 × 10^−1^. Regions of the parameter space that lead to population extinction are shaded in blue, while regions that lead to population survival are shaded green. The parameter *A* is identifiable since 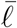 is sufficiently curved that the 95% confidence interval is relatively narrow, 2.45 × 10^−1^ ≤ *A* ≤ 5.55 × 10^−1^. To explore how these estimates lead to various long-term predictions, we randomly sample *S* = 500 parameter combinations from within the confidence interval. For each parameter sample we use the continuum model to solve for *N* (*t*), noting that some of these trajectories lead to extinction whereas others lead to survival. The lower panel of Figure 7(c) shows the prediction interval, with trajectories leading to population survival shaded green and trajectories leading to extinction shaded blue. While the MLE solution leads to extinction, many parameters from within the 95% confidence interval match early-time data reasonably well but produce incorrect predictions of population survival at late times. Overall, while the parameter *A* is reasonably well-identified by early time data, we see that some parameter combinations within the confidence interval lead to erroneous predictions of population survival at late time. This outcome highlights an important point that identifiable parameter estimates can lead to incorrect extinction risk predictions.

**Figure 7.**
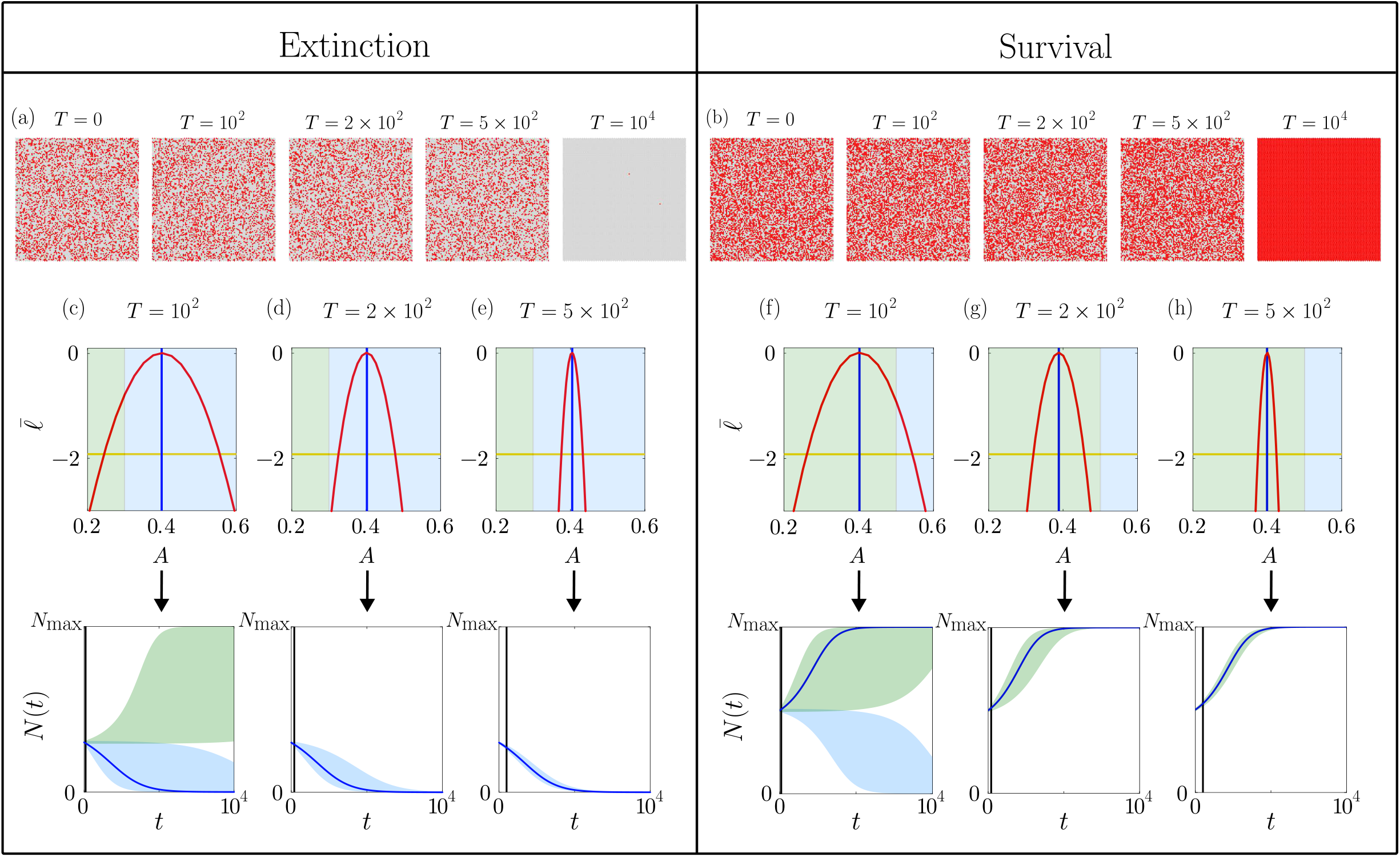
Predictions for the well-mixed initial distribution: (a)–(b) Snapshots of the discrete simulation with *B* = 0.3 for extinction, and *B* = 0.5 for survival. In (c)–(h), the normalised log-likelihood (solid red) for data collected at different observation times, *T* (solid black), parameter values associated with extinction shown in blue, whereas parameters associated with survival are shown in green. Simulation parameters are *M* = 1, *P* = 2.5 × 10^−3^, *A* = 4 × 10^−1^, *r*^(*m*)^ = 1, *r*^(*p*)^ = 4 and *N*_max_ = *IJ* = 1.16 × 10^4^.

Figures 7(d)–(e) present 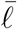 and corresponding prediction intervals for data collected at later times, *T* = 2 × 10^2^ and 5 × 10^2^. The resulting MLEs are 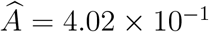 and 4.04 × 10^−1^, respectively, which closely matches the reference parameter value *A* = 4×10^−1^. In both cases the 95% confidence intervals narrows to 3.25 × 10^−1^ ≤ *A* ≤ 4.78 × 10^−1^ for data collected at *T* = 2 × 10^2^, and 3.76 × 10^−1^ ≤ *A* ≤ 4.33 × 10^−1^ for data collected at *T* = 5 × 10^2^, indicating increased inferential precision. The prediction intervals associated with this later time data lead to accurate long-time predictions where all trajectories predict long-time extinction, which is consistent with the original discrete simulations in Figure 7(a).

Additional results in Figure 7(b) show a series of snapshots of a discrete simulation with *A* = 4 × 10^−1^ in which the population eventually survives and grows to completely fill the lattice. Using count data at *T* = 1 × 10^2^ we obtain 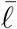 shown in Figure 7(f) where we see that the MLE, 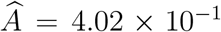, is very close to the expected result of *A* = 4 × 10^−1^. As in the extinction scenario, the parameter appears to be well identified by the data since 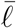 is sufficiently curved that the 95% confidence interval is reasonably well constrained, 2.61 × 10^−1^ ≤ *A* ≤ 5.43 × 10^−1^. The associated prediction interval in the lower panel in Figure 7(f) shows that some portions of the prediction interval predict long-time survival, as expected, however other parts of the prediction interval incorrectly predict population extinction. Further results in terms of 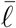 and the associated prediction intervals are given in Figure 7(g)–(h) for data collected at later times, *T* = 2 ×10^2^, and *T* = 5 ×10^2^. In these cases the MLEs are 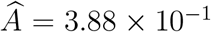 and 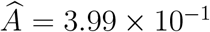. Here we see that the 95% confidence interval narrows to 3.2 × 10^−1^ ≤ *A* ≤ 4.57 × 10^−1^ for data collected at *T* = 2 × 10^2^, and 3.74 × 10^−1^ ≤ *A* ≤ 4.24 × 10^−1^ for data collected at *T* = 5 × 10^2^. The prediction intervals associated with this later time data lead to accurate long-time predictions where all trajectories predict long-time survival.

Comparing the log-likelihood functions and prediction intervals in Figure 7(c)–(e) and Figure 7(f)–(h) indicates that our ability to accurately forecast long-time survival or extinction is very sensitive to the time at which data are collected, regardless of whether we are working with a well-mixed population that eventually survives or becomes extinct.

### 6.2 One-dimensional column initial distribution

We now examine whether the conclusions from the well-mixed case are affected by the spatial arrangement of agents. This is an interesting question since Li et al. [23] demonstrated that, beyond the Allee threshold, spatial structure plays an important role in determining population survival or extinction. Unlike the well-mixed case where spatial gradients vanish, the dynamics of the 1D column distribution depend on the horizontal spatial coordinate *x*. Since the evolution of the population is driven both by 1D macroscopic spreading, proliferation and death, we will estimate both parameters, ***θ*** = (*D, A*)^⊤^. We take the parameter space to be in the range 1 × 10^−2^ ≤ *A* ≤ 8 × 10^−1^ and 2 × 10^−1^ ≤ *D* ≤ 3 × 10^−1^, our results (below) confirm that this choice of parameter space is appropriate.

Figure 8(a) presents snapshots of discrete simulations in which the population goes extinct. Figure 8(c)–(d) shows the (*D, A*)^⊤^ parameter space superimposed with a contour at 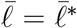 that is based on count data at *T* = 2×10^2^ and *T* = 4×10^2^, respectively. The resulting MLEs are 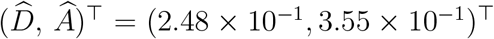 and 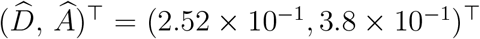, both of which are close to the expected result, (*D, A*)^⊤^ = (2.5 × 10^−1^, 4 × 10^−1^)^⊤^. In both cases the parameter confidence set is constrained to a small region indicating that the parameters are practically identifiable [26, 27]. While this early-time data identifies the parameters, the corresponding prediction intervals involve both extinction and survival outcomes, despite the initial simulated system ultimately becoming extinct. This behaviour is consistent with the well-mixed case. By *T* = 1 × 10^3^, the MLE is 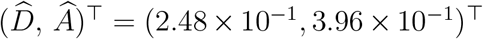 and the extent of the parameter confidence set contracts, leading to improved prediction accuracy which correctly rules out survival and predicts only extinction.

**Figure 8.**
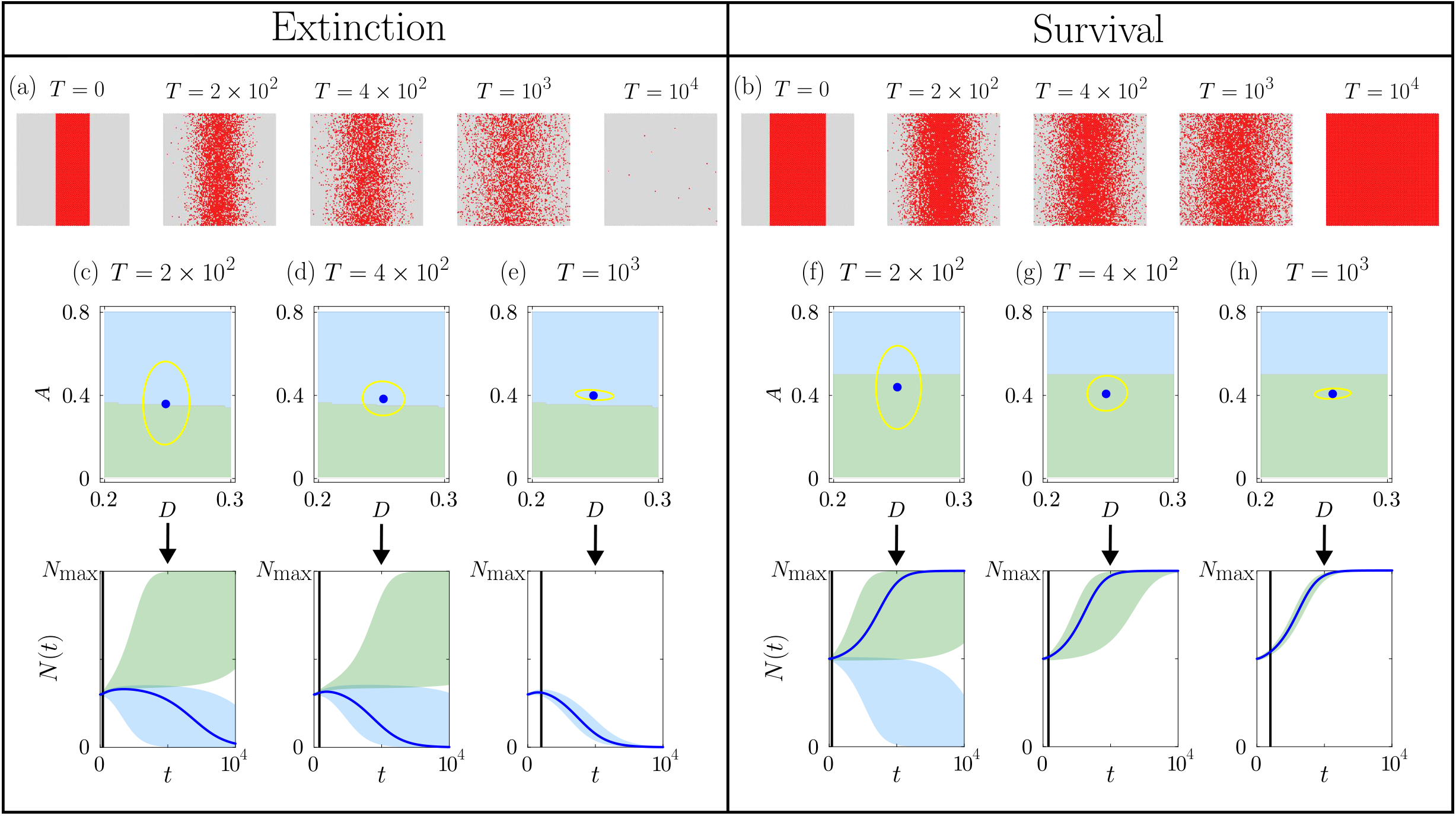
Predictions for the 1D column initial distribution: (a)–(b) Snapshots of the discrete model with *w*_1_ = 30 leading to extinction, and *w*_1_ = 50 leading to survival, respectively. (c)–(h) Top row shows the (*A, D*) parameter space with regions where the continuum model predicts extinction (blue) and survival (green). The parameter space is superimposed with contour at 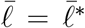 (gold curve) which encloses the 95% parameter confidence set and the MLE (blue dot). The bottom row shows the corresponding prediction interval for *N* (*t*) that includes both extinction (blue) and survival (green) predictions together with the MLE solution (blue curve). Simulation parameters are *M* = 1, *P* = 2.5×10^−3^, *A* = 4 ×10^−1^, *r*^(*m*)^ = 1, *r*^(*p*)^ = 4 and *N*_max_ = *IJ* = 1.16 × 10^4^.

Figure 8(b) shows discrete snapshots where the population survives. Using quadrat count data collected at *T* = 2 × 10^2^, 4 × 10^2^, and 1 × 10^3^, we construct parameter confidence sets and prediction intervals. As shown in Figure 8(f), the MLE at *T* = 2 × 10^2^ is 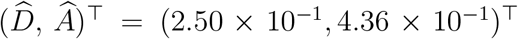. While the parameter confidence set is identifiable, the resulting prediction intervals encompasses both extinction and survival outcomes despite the population ultimately surviving. Figure 8(g)–(h) correspond to count data at *T* = 4 × 10^2^ and *T* = 1 × 10^3^, giving 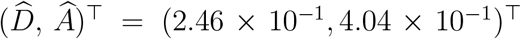 and 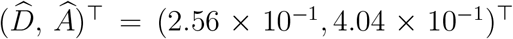, respectively. By these times, the confidence set contracts sufficiently to rule out incorrect extinction predictions.

To provide more information about the failure of early-time data to predict long-term outcomes, we select two parameter choices with 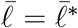 using data collected at *T* = 2 × 10^2^, as illustrated in Figure 9(a). For both parameter choices, we compute 95% prediction intervals for *N*_*u*,1_(*t*) and compare them with the observed data. Figure 9(b) shows that both parameter choices give indistinguishable prediction intervals that closely match the count data at *T* = 2 × 10^2^. The prediction intervals look almost identical because at relatively early observation times the population dynamics is primarily driven by *D* and the influence of *A* is more subtle. While the parameters are identifiable, the underlying bistability is associated with a separatrix [23, 51] on the parameter space which divides the parameter space into two regions with fundamentally distinct long-term qualitative behaviours, one leading to population survival and the other to total extinction. This means that even vanishingly small errors in parameter estimation can lead to completely different qualitative fates. As shown in Figure 9(c), when these same sample parameter sets are used to create prediction intervals at *T* = 5 × 10^3^, one parameter set incorrectly predicts survival, while the other correctly predicts extinction as evidenced by the stochastic data. This discrepancy demonstrates that while relatively early-time data may lead to practically identifiable parameter estimates, it lacks the information density required to accurately forecast the long-term qualitative fate of the population.

**Figure 9.**
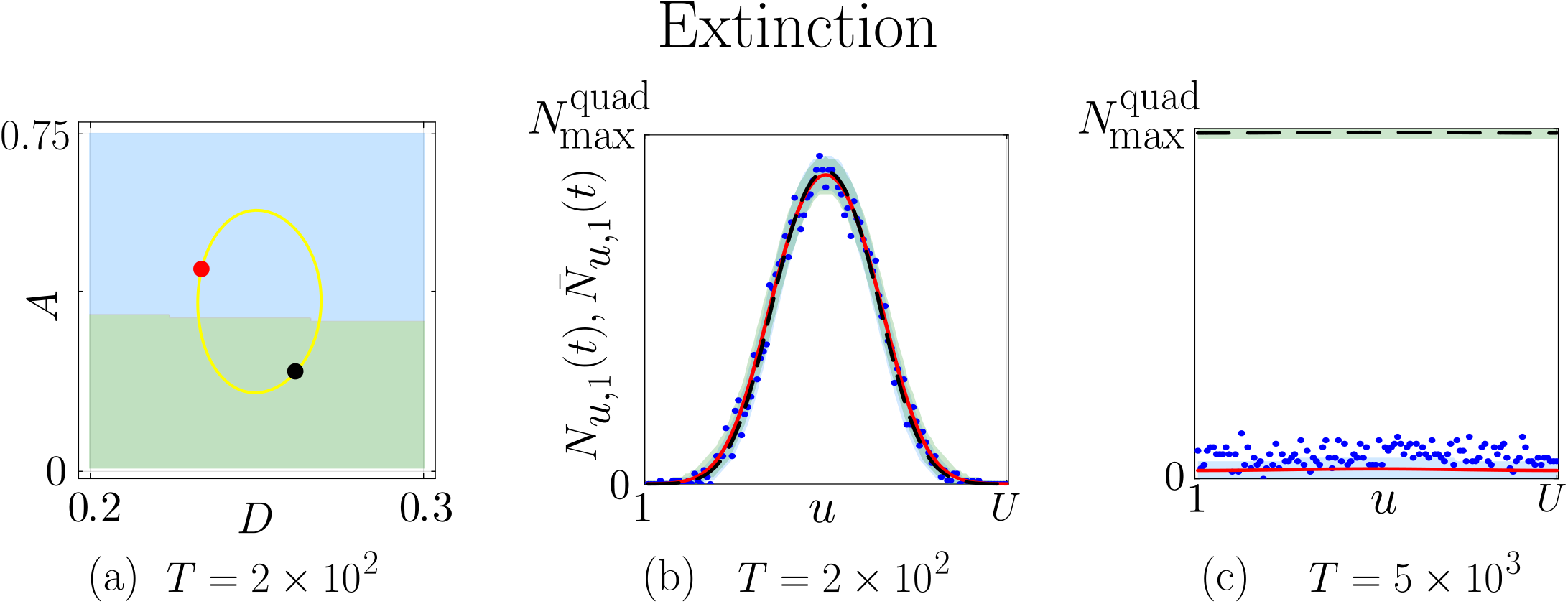
Predictions for the 1D column initial distribution: We take two points in the parameter confidence set with 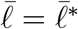: (i) (*D, A*)^⊤^ = (2.33 × 10^−1^, 4.47 × 10^−1^)^⊤^ (red dot), and (ii) (*D, A*)^⊤^ = (2.61 × 10^−1^, 2.19 × 10^−1^)^⊤^ (black dot) for data at *T* = 200. (b) Shows the associated prediction *N*_*u*,1_(2 × 10^2^) (red and black lines) together with the associated 95% prediction interval and the count data (blue dots). (c) Shows the associated *N*_*u*,1_(5 × 10^3^) (red and black lines) together with the associated 95% prediction intervals. Simulation parameters are *M* = 1, *P* = 2.5 × 10^−3^, *A* = 4 × 10^−1^, *r*^(*m*)^ = 1, *r*^(*p*)^ = 4 and *N*_max_ = *J* = 1 × 10^2^.

### 6.3 Two-dimensional square and circle initial distributions

We now consider 2D square and 2D circular initial agent distributions. The dynamics of the 2D initial conditions depends on the horizontal and vertical location, and the population dynamics is driven both by macroscopic spreading and proliferation/death, so we estimate ***θ*** = (*D, A*)^⊤^. We take the parameter space to be in the range, 1 × 10^−2^ ≤ *A* ≤ 1 and 2 × 10^−1^ ≤ *D* ≤ 3 × 10^−1^, our results (below) confirm that this choice of parameter space is appropriate.

Results in Figure 10 consider the 2D square initial distribution of agents. Figure 10(a) presents snapshots of discrete simulations in which the population ultimately goes extinct. Figures 10(c)–(e) show the (*D, A*)^⊤^ parameter space superimposed with a contour at 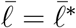 using data at *T* = 2 × 10^2^, *T* = 4 × 10^2^, and *T* = 1 × 10^3^. These results give 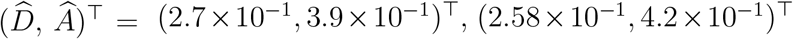, and (2.48 ×10^−1^, 4 ×10^−1^)^⊤^, respectively. Again, the 95% confidence set constructed using early-time data is confined to a relatively small region of the parameter space, and the size of the confidence set contracts significantly when constructed using count data collected at later observation times. Prediction intervals constructed from parameters estimated using count data collected at *T* = 2 × 10^2^ incorrectly predict both survival and extinction outcomes, whereas prediction intervals constructed from parameters estimated using count data collected at *T* = 4×10^2^ and *T* = 10^3^ correctly predict extinction.

**Figure 10.**
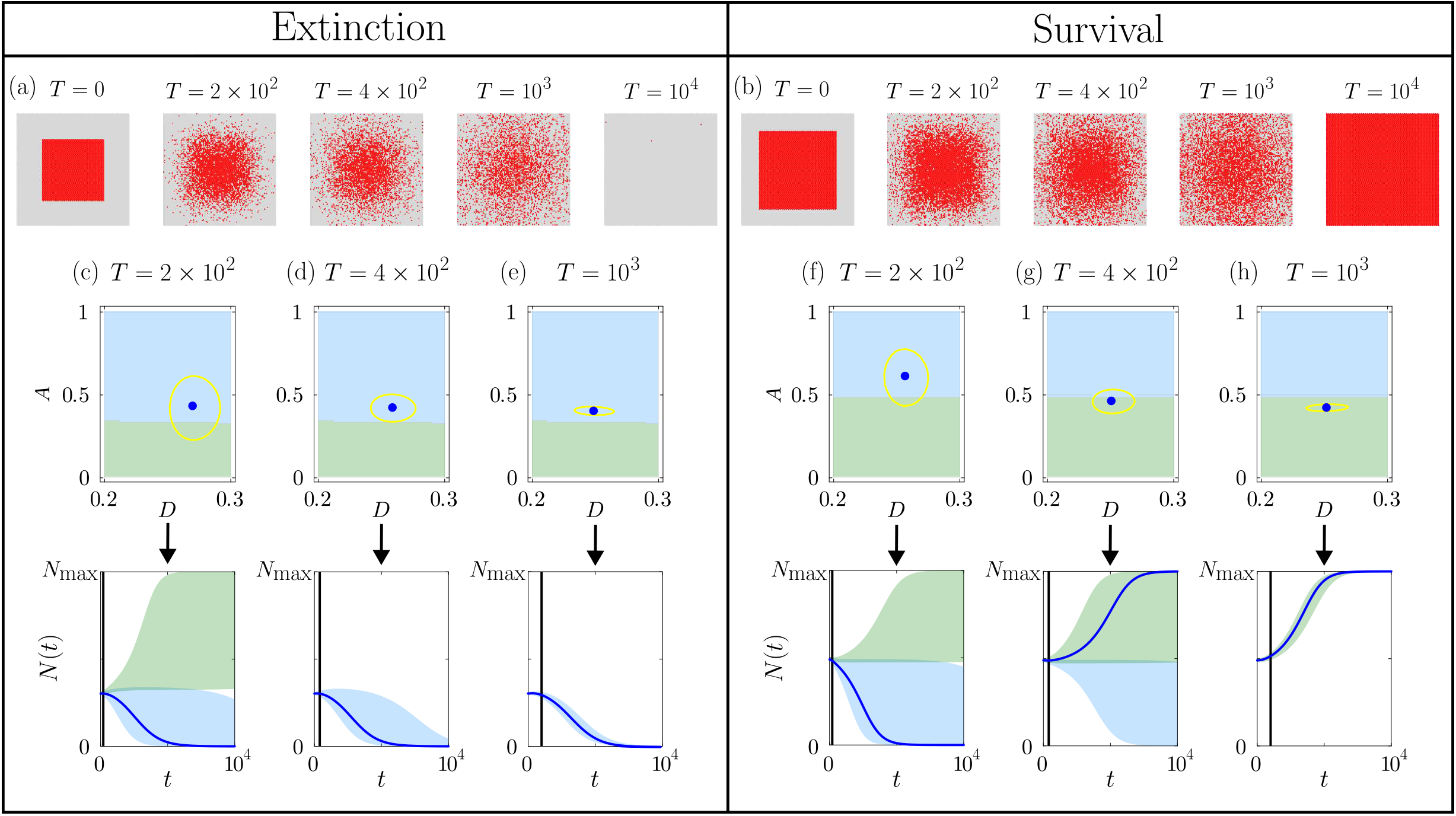
Predictions for the 2D square initial distribution: (a)–(b) Snapshots of the discrete models with *w*_2_ = 55 leading to extinction, and *w*_2_ = 70 leading to survival, respectively. (c)–(h) Top row shows the (*A, D*) parameter space with regions where the continuum model predicts extinction (blue) and survival (green). The parameter space is superimposed with contour at 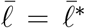 (gold curve) which encloses the 95% parameter confidence set and the MLE (blue dot). The bottom row shows the corresponding prediction interval for *N* (*t*) that includes both extinction (blue) and survival (green) predictions together with the MLE solution (blue curve). Simulation parameters are *M* = 1, *P* = 2.5×10^−3^, *A* = 4 ×10^−1^, *r*^(*m*)^ = 1, *r*^(*p*)^ = 4 and *N*_max_ = *IJ* = 1.16 × 10^4^.

Figure 10(b) shows snapshots of a discrete simulation where the population ultimately survives. Figures 10(f)–(g) show the (*D, A*)^⊤^ parameter space superimposed with a contour at 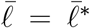 based on count data collected at *T* = 2 × 10^2^ and *T* = 4 × 10^2^. These results give 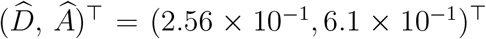 and 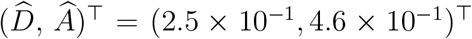, respectively. The parameter confidence set based on early time data cannot rule out longer-term survival or prediction. In this case it is notable that the MLE parameter set based on early time data incorrectly predicts extinction. Data collected at *T* = 1 × 10^3^ leads to 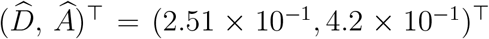 which is close to the reference value. For data collected at this later observation time, the confidence set has contracted further, and incorrect extinction predictions are no longer observed.

Figure 11 shows snapshots using 2D circular initial distribution of agents, where results are qualitatively similar those for the 2D square configuration presented in Figure 10. Snapshots of a discrete simulation undergoing extinction is illustrated in Figure 11(a). Figures 11(c)– (e) show the (*D, A*)^⊤^ parameter space superimposed with a contour at 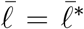 using data collected at *T* = 2×10^2^, 4×10^2^, and 1×10^3^. These give 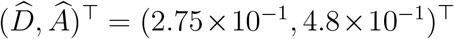, (2.75×10^−1^, 4.3×10^−1^)^⊤^, and (2.58×10^−1^, 4.1×10^−1^)^⊤^, respectively. The 95% confidence set constructed from early-time data confines a relatively small region of the parameter space, and the size of the confidence set contracts significantly when constructed using count data collected at later observation times. Prediction intervals constructed from early time data incorrectly predict both survival and extinction, whereas prediction intervals constructed from later times correctly predict extinction only.

**Figure 11.**
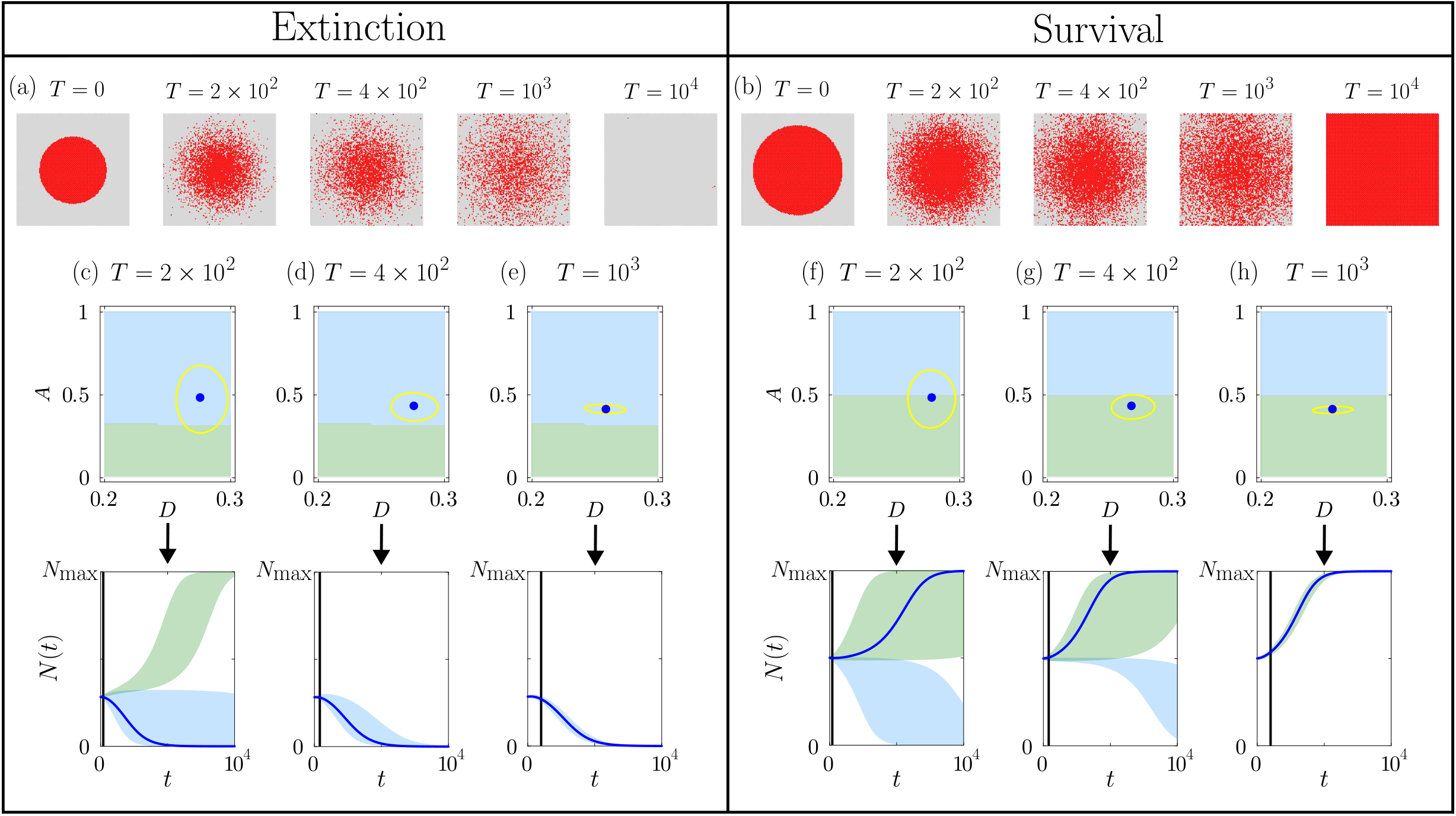
Predictions for the 2D circular initial distribution: (a)–(b) Snapshots of the discrete models with *w*_3_ = 60 leading to extinction, and *w*_3_ = 80 leading to survival, respectively. (c)–(h) Top row shows the (*A, D*) parameter space with regions where the continuum model predicts extinction (blue) and survival (green). The parameter space is superimposed with contour at 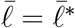 (gold curve) which encloses the 95% parameter confidence set and the MLE (blue dot). The bottom row shows the corresponding prediction interval for *N* (*t*) that includes both extinction (blue) and survival (green) predictions together with the MLE solution (blue curve). Simulation parameters are *M* = 1, *P* = 2.5 × 10^−3^, *A* = 4 × 10^−1^, *r*^(*m*)^ = 1, *r*^(*p*)^ = 4 and *N*_max_ = *IJ* = 1.16 × 10^4^.

Figure 11(b) shows snapshots from a discrete simulation where the population survives. Figures 11(f)–(g) show the (*D, A*)^⊤^ parameter space superimposed with a gold contour at 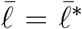 using data collected at *T* = 2 × 10^2^ and 4 × 10^2^. This leads to 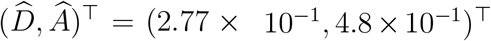 and (2.66 × 10^−1^, 4.3 × 10^−1^)^⊤^ with the prediction intervals spanning both survival and extinction regimes. By *T* = 1×10^3^, the confidence interval contracts sufficiently to provide accurate survival predictions with 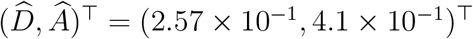.

## 7 Conclusion and Future Work

In this work we introduce an efficient, practically-motivated likelihood-based workflow using quadrat-based count data for identifiability, estimation and prediction for bistable populations. The workflow enables systematic estimation of model parameters and the construction of parameter confidence sets and prediction intervals, allowing us to directly relate parameter uncertainty to qualitative outcomes such as population survival or extinction. This is achieved without relying on expensive simulation-based inference methods. The workflow is validated by generating noisy count data from the discrete model described by Li et al. [23] where agent proliferation is governed by a crowding function *F* (*K*) = (1*/A*)(1 − *K*)(*K* − *A*), which induces a strong Allee effect with a threshold *A*. This formulation permits either population survival or extinction depending on the initial population size, spatial configuration, diffusivity *D*, and the Allee threshold *A*. Numerical solutions of the continuum limit PDE for this discrete model accurately describes the mean behaviour of the discrete count data (Section 4), thereby providing a benchmark against which accuracy of the inferred parameter estimates can be assessed.

Our results demonstrate that accurate prediction of survival or extinction depends critically on the quality, quantity, and spatial resolution of the available data. In particular, parameter estimates may appear identifiable while still leading to incorrect predictions, as observed with early-time count data. We find that *D* can be estimated reliably from relatively early-time observations, whereas *A* has a more subtle influence. At intermediate–to-late times, the influence of *A* becomes more pronounced, reducing the range of plausible parameter values and leading to improved parameter estimation and more reliable long-term predictions. Beyond the timing of data collection, the initial spatial arrangement of the population plays a role in determining parameters that ultimately distinguish between survival and extinction scenarios. This is consistent with previous studies demonstrating that spatial population structure can influence survival and extinction outcomes [44].

A limitation of the present study is that we restrict our attention to estimating either one or two parameters, ***θ*** = (*D, A*)^⊤^, while keeping *C*(*x, y*, 0) and *λ* fixed. In this work we made this decision for the sake of simplicity and transparency; however, the same workflow applies to higher-dimensional parameter spaces [25]. In these cases greater care needs to be taken in terms of interpreting and virtualizing 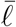, potentially using the profile likelihood to visualise the log-likelihood function when more than two unknown parameters are of interest [24, 26].

Throughout this study we use count data collected at a single time point. Incorporating data at several time points is straightforward to incorporate into the workflow, and may lead to reduced uncertainty in parameter estimates, more restricted parameter confidence sets, and accurate survival or extinction predictions. While we restricted our investigation to agent motility corresponding to linear diffusion, the underlying discrete model can be extended to incorporate non-linear diffusion by using different movement crowding functions [44]. Applying the workflow to estimate model parameters that include more complicated motility mechanisms corresponding to non-linear diffusion is an interesting avenue for future research where the same workflow steps will apply. Finally, while this simulation study focuses on synthetic data generated from a known discrete model, our workflow can also be used to interpret real-world datasets [25], and we leave this task for future consideration.

## Data Accessibility

Julia implementations within Jupyter notebooks for all computations are available on GitHub at https://github.com/Adarsh-KrishnanRajkumar/Rajakumar2026_DataRequirements_AlleeInference.

## APPENDIX A Numerical Methods

The continuum limit description for the well-mixed initial distribution is given by

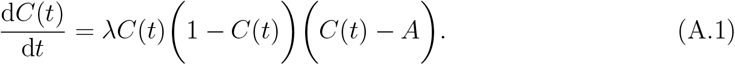

We solve this ordinary differential equation (ODE) numerically using computational methods described later in this Appendix.

The solution of the continuum limit description for the column initial distribution (Equation (1)) is obtained by discretising the domain 0 *< x < W* using a uniform mesh with mesh spacing *δ*. Each mesh point has a position of *x*_*i*_ for *i* = 1, 2, 3, …, I and we write *C*(*x*_*i*_, *t*) as *C*_*i*_(*t*). Discretising spatial derivatives using the standard central difference approach leads to

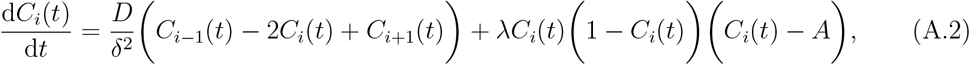

for *i* = 2, 3, …, ℐ − 1. We set *C*_1_ = *C*_ℐ_ at the boundaries to simulate periodic boundary conditions. This system of ODEs are solved numerically using computational methods described later in this Appendix. For this numerical approximation we take *δ* = *W/*300, and additional results (data not shown) indicate that this choice leads to grid-independent results for the problems that we consider.

The solution of the continuum limit description for the 2D initial conditions (Equation (1)) is obtained by discretising 0 *< x < W* and 0 *< y < H* uniformly with mesh spacing *δ*_*x*_ = *W/*100 and *δ*_*y*_ = *H/*100. Each mesh point has a position of (*x*_*i*_, *y*_*j*_) for *i* = 1, 2, 3, …, I and *j* = 1, 2, 3, …, J and we write *C*(*x*_*i*_, *y*_*j*_, *t*) as *C*_*i, j*_(*t*). Discretising spatial derivatives using the standard central difference approach leads to

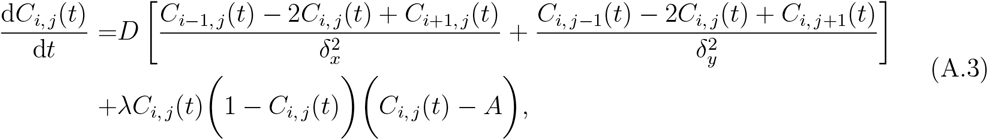

which is valid for *i* = 2, 3, …, ℐ − 1 and *j* = 2, 3, …, J − 1. The discretised equations on boundary nodes are adjusted to simulate periodic boundary conditions. We solve this system of ODEs numerically using computational methods described later in this Appendix.

Additional results (not shown) indicate that our choice of *δ*_*x*_ and *δ*_*y*_ leads to grid-independent results for the problems that we consider.

For all three spatial configurations, the resulting ODEs are integrated through time using the `DifferentialEquations.jl` library in Julia [52]. For simplicity, we use Heun’s method with automatic time-stepping and standard absolute and relative error tolerances.

